# Pathological α-Synuclein Perturbs Nuclear Integrity

**DOI:** 10.1101/2025.03.04.641384

**Authors:** Michael Millett, Allison Comite, Elisabeth Martin Castosa, Anika Heuberger, Preston Wagner, Ignacio Gallardo, Jessica Reinhardt, Nicole Chambers, Dominic Hall, Evan Miller, Douglas Nabert, Barbara Balsamo, Stefan Prokop, Mark Moehle

## Abstract

Pathological aggregates of α-synuclein are a hallmark of a group of neurodegenerative disorders collectively termed synucleinopathies. The physiological function of α-synuclein, and the detrimental effects of the pathological variants of α-synuclein have been widely debated, but recent evidence has suggested an emerging consensus on a critical role for α-synuclein in regulating synaptic function. However, a controversial role for α-synuclein in nuclear function in both normal and pathogenic states has been proposed, and the degree to which α-synuclein localizes within the nucleus and subsequent impact on the nucleus are poorly understood. To begin to address this controversy, we employed synucleinopathy murine and cell culture models, as well as postmortem human Lewy Body Dementia tissue to elucidate the extent to which pathological α-synuclein localizes within the nuclear compartments, and the downstream consequences of this localization. We observed pathological aggregation of α-synuclein within the nucleus in both murine models and human postmortem Lewy Body Dementia cortex via quantitative super resolution microscopy. In both mouse and human brain tissue the presence of α-synuclein in the nucleus correlated with abnormal morphology of nuclei. This pathological accumulation of α-synuclein in the nucleus was not observed in control mice, human tissue without pathology, or control cells. We subsequently examined the mechanistic consequences of pathological accumulation of α-synuclein in the nucleus. Synucleinopathy models displayed increased levels of the DNA damage marker 53BP1. Furthermore, cells with pathological α-synuclein exhibited elevated markers of nuclear envelope damage and abnormal expression of nuclear envelope repair markers. Our cell culture data also suggests altered RNA localization in response to pathological α-synuclein accumulation within the nucleus. Lastly, we show that nuclear Lewy-like pathology leads to increased sensitivity to nuclear targeted toxins. Taken together, these results rigorously illustrate nuclear localization of pathological α-synuclein with super resolution methodology and provide novel insight into the ensuing impact on nuclear integrity and function.

## Introduction

Alpha-synuclein (α-syn) is a small, 140 amino acid protein, that is predominantly expressed in neurons, compromising up to 1% of the total protein content of the brain [32]. Pathological α-syn (p-α-syn) is a hallmark feature of a group of neurodegenerative diseases, termed synucleinopathies, in which α-syn accumulates into hyperphosphorylated, ubiquitinated, insoluble, and high molecular weight aggregates. Pathological α-syn aggregates are found throughout multiple cell types and brain regions across synucleinopathies. α-Syn was initially identified in the electric lobe of the *Torpedo californica*, with expression spanning from the presynaptic terminal to within the nucleus [32, 39]. In decades of research since its initial discovery, α-syn has been shown to have a critical role in synaptic function such as neurotransmitter release and regulation of synaptic vesicle dynamics in mammals in both pathological states and normal physiology [9, 24, 30, 54]. However, a role for α-syn in nuclear function has remained controversial.

There are a number of reports of both homeostatic and pathogenic α-syn variants having nuclear localization and function in models of synucleinopathies (McLean et al., 2000; Mori et al., 2002; Schaser et al., 2019). Recent studies have indicated that α-syn may directly bind DNA, and have a role in DNA damage repair [52]. Overexpression of α-syn within the nucleus is strongly correlated with nuclear stress, nuclear dysfunction, altered behavioral outcomes, and increased neurodegeneration [12, 43]. However, studies that rely on over-expression of α-syn are obscured by the fact that small proteins less than 40 kDa can passively diffuse into the nucleus [23].Furthermore, pathogenic *SCNA* mutations have been shown to impact nuclear envelope integrity and function [10]. Additionally, in postmortem studies of human tissue, both normal and pathological α-syn, termed Lewy pathology [27], have been suggested within the nucleus via immunofluorescence microscopy and biochemical assays [25]. Despite this body of evidence, the function or localization of α-syn in the nucleus has remained controversial due to insufficient resolution to definitively determine localization, and a number of initial reports being confounded by off target binding of initial α-syn antibodies in the nucleus [20, 25].

Here, we address the nuclear localization of p-α-syn utilizing the pre-formed fibril (PFF) model of α-synucleinopathies in rodents and cell culture, and confirm many of our findings in post-mortem cortical brain tissue from patients with synucleinopathies. We characterize the extent to which pathological α-syn localizes to the nucleus via quantitative super-resolution microscopy. Additionally, we elucidate the effects of nuclear localization on envelope integrity, nucleic acid damage, and function of the nucleus with biochemical and cell-based analysis. Our results identified nuclear envelope abnormalities, DNA damage, increased susceptibility to nuclear insult, and potential deficits in envelope repair resulting from the nuclear and peri-nuclear localization of pathological α-syn. Nuclear p-α-syn and aberrant nuclear morphology was confirmed with postmortem human tissue. Our data provides additional insight into nuclear α-syn in the context of synucleinopathies and suggests new mechanisms to the etiology of these neurodegenerative disorders.

## Materials and Methods

### α-Syn Monomer Production

Glycerol stocks of BLD21 E.coli cells transfected with PRK172-α-synuclein plasmid (kind gift of Dr. Michael Goedert) were removed from storage and was transferred into a starter flask containing Terrific Broth (VWR) supplemented with 100 μg/mL ampicillin (VWR) and shaken overnight at 37 °C. The next day, starter culture was transferred to large culture flasks and incubated further until OD_600_= 0.8, at which point synuclein expression was induced with IPTG for 3 hours. Cell pellets were collected, lysed through sonication, boiled at 100C, and then centrifuged. The supernatant was collected then dialyzed overnight in SnakeSkin Dialysis Tubing, 3.5 MWCO (ThermoFisher) with gentle stirring in dialysis buffer containing 10mM Tris, 100mM NaCl, 1mM EDTA, 1mM PMSF, pH =7.6. The dialyzed lysate was then loaded onto a HiPrep Q HP 16/10 anion exchange chromatography (AEC) column (Cytiva) via a BioRad NGC Quest FPLC. After binding, α-synuclein was eluted from the column through a gradient of low salt buffer “A” (10mM Tris, 50 mM NaCl, 1mM EDTA, ph7.6) and high salt buffer B (10 mM Tris, 1M NaCl, 1mM EDTA, pH7.6) from 0% to 60% buffer B over 120 minutes at a flow rate of 1.0 mL/minute. Elution of α-synuclein typically occurs at ∼30% buffer B. Probable fractions containing monomeric α-synuclein determined by OD_280_ nm peaks on the chromatograph, were loaded on 4-20% precast Tris-Glycine gels (Bio-Rad). The loaded gels were run at 120V to completion, briefly washed, and subsequently stained with GelCode Blue (ThermoFisher). α-syn-positive fractions (Sup. Fig. 1) were dialyzed overnight as described above at 4°C in dialysis buffer II (10mM Tris pH =7.6, 50mM NaCl). The following day, sample was subjected to size exclusion chromatography (SEC) via running on a HiLoad 16/600 Superdex 200pg SEC column (Cytiva). Probable fractions were selected and processed as above (Sup. Fig. 1), before concentration to 10 mg/mL via centrifugation in Ultra-15 centrifugal filters (Amicon). Concentrated α-syn was incubated on High-Capacity Endotoxin Removal Resin (Pierce) for 1 hour to remove trace endotoxin as per kit instructions. The endotoxin-free monomeric synuclein was then concentrated to 12 mg/ml and stored in 50mM Tris-HCl, 150mM KCl, pH7.5 buffer at -80°C until PFF production.

### PFF Production

PFFs were manufactured as previously described in literature (Patterson et al., 2019). Briefly, monomeric alpha-synuclein was thawed on ice and centrifuged 10 minutes at 15,000 xg at 4 °C. Supernatant was removed, any pellet was discarded. Concentration of thawed monomeric alpha-synuclein was determined via a spectrophotometer using Beers Law and diluted to 5 mg/ml in 50mM Tris-HCl, 150mM KCl, pH7.5 buffer in low-binding microcentrifuge tubes (Eppendorf). Tubes were placed into a thermal mixer (Eppendorf) inside a 37 °C oven and left to shake for 7 days at 1000 rpm. After shaking, the sample was briefly centrifuged, supernatant discarded, and pellet resuspended in half of the initial shaking volume of fibrilization buffer (50mM Tris-HCl, 150mM KCl, pH7.5). A small aliquot of PFFs was mixed at an equal ratio with 8M guanidinium chloride (GnCl) and absorbance was measured on a NanoDrop (ThermoFisher) spectrophotometer. Beers law was used to determine PFF concentration from absorbance and sample was diluted to 5 mg/mL for short term storage and/or subsequent sonication.

### Sonication and Dynamic Light Scattering

PFFs were transferred to 50 µl microtubes (Covaris) and placed into the sample holder of a M220 Focused-ultrasonicator (Covaris). Sonication protocol was executed with the following parameters: peak incidence power of 50%, duty factor of 10%, 1 second treatment time, 200 cycles per burst, and 220 iterations. Temperature control was set to 12°C +/-2°C. Proper sonication was verified via dynamic light scattering (DLS) analysis. A sonicated PFF aliquot was diluted 1:4 with PBS and transferred to a micro-capillary (Bettersize) via capillary action. Capillaries were placed into a BeNano 180 Zeta Pro (Bettersize) machine and the sizing protocol was performed. The average target size of 50 nm was used as quality cutoff (Sup. Fig. 1), and sample > 10 nm above the cutoff were either rejected or re-sonicated to specification.

### Surgery

Male C57BL/6J mice 10-14 weeks of age were anesthetized with 1-3% isoflurane in an induction chamber, and then transferred to a nose cone of an RWD or Stoelting stereotaxic frame. Mice were given Meloxicam (15 mg/kg, s.c.) as a preoperative analgesic, and Bupivacaine (0.2-0.3 mL at a dose of 5 mg/mL, S.C.) as a local analgesic at the site of incision. Fur was removed from the incision site via shaving prior to stabilization in a stereotaxic apparatus. Mice remained under anesthesia with a constant 1-1.5% isoflurane at between 1-2 L/min flow. The surgical site was disinfected with 4% chlorhexidine solution and sterile saline solution using cotton-tipped applicators. Burr holes were drilled above bilateral target coordinates (AP: 0.2, ML: +/-2.0, DV: -2.8 from dura matter) using an Micro-Drill (IDEAL Part No. 67-1204). A 10 µL micro-syringe (Hamilton 1701 RN) was used to inject 2 µL of PFF or control at a rate of 0.5 µL/min/site. Following the injection, the needle was left in place for 5 minutes to allow for absorption before being slowly withdrawn. The surgical site was sutured using nylon 5-0 surgical sutures and fixed in place using biocompatible glue (Vetbond) directly on the sutures. Mice were given 1 mL of saline at the end of the surgery to stave off dehydration, and were monitored postoperatively for two days after surgery, during which Meloxicam was given daily (15 mg/kg, S.C.).

### Tissue Processing

At the desired time points post PFF injection, mice were anesthetized with isoflurane and transcardially perfused with ice-cold PBS followed by 4% paraformaldehyde in PBS. Brains were carefully extracted from the skull and post-fixed in 4% PFA in PBS for 24 hours before short-term storage in PBS with 0.05% sodium azide. Whole brain tissue was mounted and processed on a Lieca Vibratome 1200S, calibrated to 18° clearance angle in ice-cold PBS. Slices 30-100 μm were obtained and stored in 24-well plates in PBS with 0.05% sodium azide at 4°C until downstream use.

### Cell Culture

Neuro 2a cells were cultured in 6-well plates in DMEM medium supplemented with 5% FBS and 1x GlutaMAX (Gibco). Cells were housed in a 37°C humidified incubator with 5% CO_2_ and ongoing cultures were subjected to media changes every 48 hours. At 70-80% confluency cells were dissociated with StemPro Accutase (Gibco) and passaged at a 1:4 ratio into new 6-well plates.

To create a population of cells with α-syn pathology, we subjected cells to incubation with α-syn PFFs. First, 5 mg/mL aliquots of PFFs were thawed and sonicated as described above. Next, PFFs were diluted 1 to 3:5 and mixed with BioPORTER QuickEase Protein Delivery Reagent (Millipore Sigma) for 5 minutes. The BioPORTER & PFF mixture was subsequently diluted up to 1.5 mL. The final concentration of PFF in serum free media was 5.72 µM. A single well of a 6-well dish was briefly washed 2x with serum free media and then incubated with the PFF/BioPORTER for 4 hours. Control wells were also subjected to serum free conditions. After 4 hours, the PFF media was aspirated, and cells were placed in normal growth medium. PFF-subjected cells were cultured separately and were incubated with PFFs as described each passage thereafter up to 6 total PFF incubations.

### Immunofluorescence

All immunofluorescence labeling of tissue was performed in 24-well plates. Slices made as above were briefly washed with PBS before 2 hours incubation in blocking/permeabilization buffer (5% normal donkey serum, 0.1% triton x-100) at room temperature with gentle-shaking. Free-floating slices were next transferred into wells with primary antibody (See Table 1) for overnight incubation at 4°C with gentle shaking. The next morning, tissue was washed 3x with PBS and stained with secondary antibodies for 1 hour at room temperature. All antibodies and their dilutions are listed in Table 1. Mouse antibodies were diluted in mouse-on-mouse (M.O.M) diluent buffer (Vector laboratories) for labeling of murine tissue. All other antibodies were diluted in blocking/permeabilization buffer for murine and human tissue. After secondary labeling, samples were subsequently washed 3x with PBS, incubated in PBS with DAPI for 5 minutes, and washed 2x more times with PBS before imaging. Labeled slices were stored at 4°C in PBS .05% sodium azide.

**Table 1:**
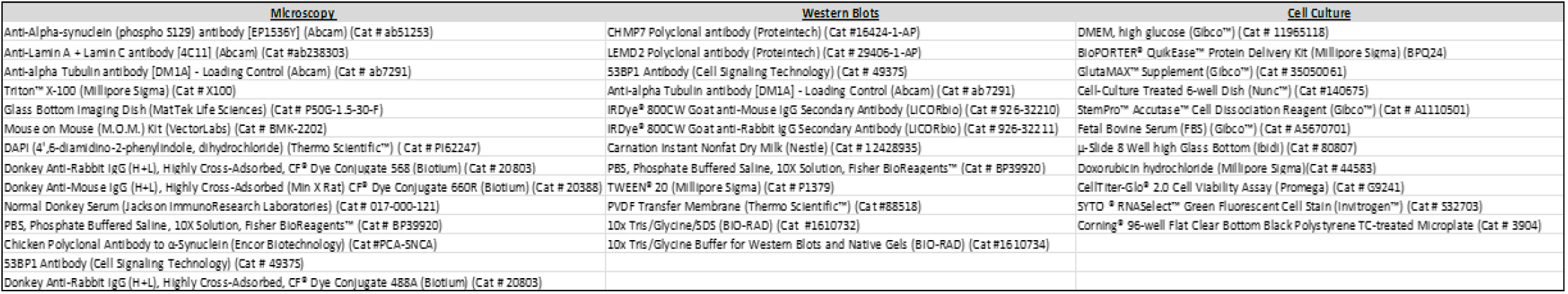
Materials and Reagents: Microscopy (Left Column), Western Blots (Center Column), and Cell Culture (Right Column).

Cell culture immunofluorescence was performed as above except Neuro 2a cells were plated into 8-well chamber slides (Cellvis) coated with Matrigel (Corning) for antibody labeling and imaging. For SYTO RNASelect labeling, the manufactures (Invitrogen) instructions were followed. Briefly, 500 nM solution was prepared in serum free media, incubated with cells for 20 minutes in a cell culture incubator. Cells were then post fixed in methanol at -20°C for 10 minutes, washed, and stored in PBS for imaging.

### Western Blot Analysis

For each biological replicate, PFF and Control samples were processed together and under identical conditions. First, Neuro 2a cells were washed 1x with PBS and subsequently incubated in StemPro™ Accutase™ for 3 minutes at 37°C to dissociate cells. An equal volume of media was added to the culture dish and pipetted gently against the dish surface to detach any remaining cells. Next, cells were collected into a 15 mL sterile tube, centrifuged at 500 xg for 5 minutes, supernatant aspirated, and cell pellets were resuspended in 2x Lamelli sample buffer (Bio-Rad). Sample was then briefly sonicated at 70% amplitude with a probe tip sonicator, heated for 10 minutes at 60°C, and loaded onto 4-20% gradient SDS-PAGE gel. Gels were run at 120V to completion and transferred to Polyvinylidene fluoride (PVDF) membranes overnight at 20V and 4°C. Membranes were then briefly washed in PBS with 0.1% tween 20 (PBS-T) before blocking in 5% milk in PBS-T for 2 hours. Membranes were then incubated in primary antibodies overnight at 4°C with gentle shaking. The following morning, membranes were washed 3x with PBS-T and then incubated with fluorescent secondary antibodies for 1 hour. All antibodies used for biochemical analysis are listed in Table 1. After 3 final PBS-T washes, fluorescent signal in the 800 nm channel was captured on a Licor Odyssey XF imager. Fluorescent signal was quantified in Image studios software (LICORbio) and normalized to α-tubulin loading controls.

### Spinning Disc Confocal Microscopy

Tissue slices were transferred to a glass bottom dish (MATTEK) with #1.5 cover glass for imaging. The correction collars on 60x water immersion objective (Nikon) were set to 0.17 and water was applied to objectives before and during imaging sessions. For CSU-W1 SoRa (Nikon) super resolution imaging, 4x magnification lens and 16-bit sensitive image acquisition settings were used. Images were acquired with 0.3 μm step size throughout the entire z-span of the cell(s) of interest. Diffraction-limited 60x objective images were acquired with 1x magnification. Saturation filters were enabled during imaging. Secondary antibody only controls were used to help determine acquisition settings and identify off target signal (Sup. Fig. 2). All acquisition for quantitative analysis was performed with identical settings between groups. Murine p-α-syn images were acquired with 561 nm laser power set to 7% and 300 ms exposure time. Lamin A/C was captured with 690 nm laser power set to 43.6% and 500 ms exposure time. Human Lewy pathology was acquired with 561 nm laser power set to 21.5% and 500 ms exposure time. Human nuclei were acquired with 405 nm laser power set to 23.3% with an exposure time of 500 ms. Three of the Lewy aggregates in human tissue were imaged on a Nikon CSU-W1 Spinning Disc, with Gataca super resolution modules and 100x silicon immersion lens. This system achieves similar resolution to SoRa super resolution imaging and our conservative “cutoff” resolution parameters (See Supplemental Figure 4) were deemed appropriate for these images as well. 53BP1 puncta were imaged in murine nuclei using 488 nm laser set to 16.5% laser power and with 500 ms exposure. SYTO RNASelect was imaged in Neuro 2a cells with 488 nm laser power set to 11.5% and an exposure time of 500 ms.

### IMARIS Image Analysis

All images were imported into IMARIS software (Oxford Instruments) for analysis and deconvolved with 50 μm pinhole size and water immersion settings selected. Images captured with 10x magnification air objectives had the air immersion setting selected. After deconvolution, the surfaces’ function was used to create physical structures for nuclear lamina, DNA, and pathological α-syn for various analysis. Surfaces were created off absolute intensity thresholding; with manual limits of 929.58-6042.27 for lamina, 251.685-868 for p-α-syn. Surface grain detail was set to 0.15 µm for lamina, and 0.1 µm for p-α-syn. Human pathology was surfaced with absolute intensity, manual thresholding of 1385.42-7707.77, and surface grain of 0.0378. DAPI was surfaced from local contrast, with manual thresholding (background subtraction) of 25.5, and surface grain of 0.095 µm. Creation of surfaces from fluorescent signal enables analysis in higher detail (Sup. Fig. 3). 53BP1 puncta were analyzed with spots function within IMARIS using an estimated diameter of 0.5 µm and quality thresholding of 635.

### Cell Viability Assay

Neuro 2a cells were seeded into opaque walled 96-well plates (Corning) and cultured overnight. Next, Neuro 2a cells were incubated with increasing doxorubicin concentrations in triplicate, ranging from 0.0078 µM-10 µM, for 24 hours in fresh cell culture medium. After doxorubicin incubation, Neuro 2a cells were subjected to the CellTiter-Glo 2.0 Assay according to the manufacturers (Promega) protocol in order to quantify cell viability differences. Briefly, a volume of CellTiter-Glo 2.0 reagent equal to cell culture media volume per well (100 µL) was added to each well, and cells were lightly shaken on an orbital shaker for 2 minutes. Cell culture plates were then equilibrated for 10 minutes at room temperature before recording luminescence on a BioTek Microplate Reader (Agilent). Signal was normalized to wells containing only media to account for background luminescence. Data was further normalized to the highest viability value recorded as 100% viability.

## Results

### Synucleinopathy Mice Display Robust Cortical Lewy-like Pathology

To characterize the morphology of pathological α-syn aggregates in murine brain tissue we employed an antibody specific to p-α-syn phosphorylated on Serine 129 (Abcam EP1563Y), an established biomarker for p-α-syn [4, 15, 42, 44]. All reagents and materials for tissue immunofluorescence can be found in Table 1.

Abundant p-α-syn signal was observed in the mouse cortex, specifically in deep cortical layers, consistent with prior literature after injection of PFFs into the dorso-lateral striatum (Fig. 1A). Super resolution imaging of PFF-injected cortical neurons revealed complex ultrastructural shapes of Lewy-like pathology. Analysis of these images revealed six characteristic nuclear or perinuclear morphologies of p-α-syn in the cortex (Fig. 1B). We defined these morphological shapes based on the following criteria. Type I is defined as a cell body filling structure that has pathological synuclein continuously extending into an axon or dendrite from the structure present in the cell body which often resembles a harp. Type II is a ring-like structure of pathological alpha-synuclein around the nucleus that has a loop connecting two sides of the ring structure into a “basket-like” structure. Type III is defined as multiple skein-like structures of alpha-synuclein that emanate from a common point that form a cage-like structure of alpha-synuclein in the cell body around the nucleus. Type IV is defined as a ring-like structure of alpha-synuclein in the cell body. These are differentiated from Type II aggregates by the lack of a connecting loop of synuclein between sides of the ring. Type V aggregates are defined as >5 puncta within the nuclear compartment that do not form connected structures. Type VI aggregates are long rod like inclusions that are not present in the cell body but are present in dendrites and axons, and are similar in appearance to Lewy neurites.

**Figure 1:**
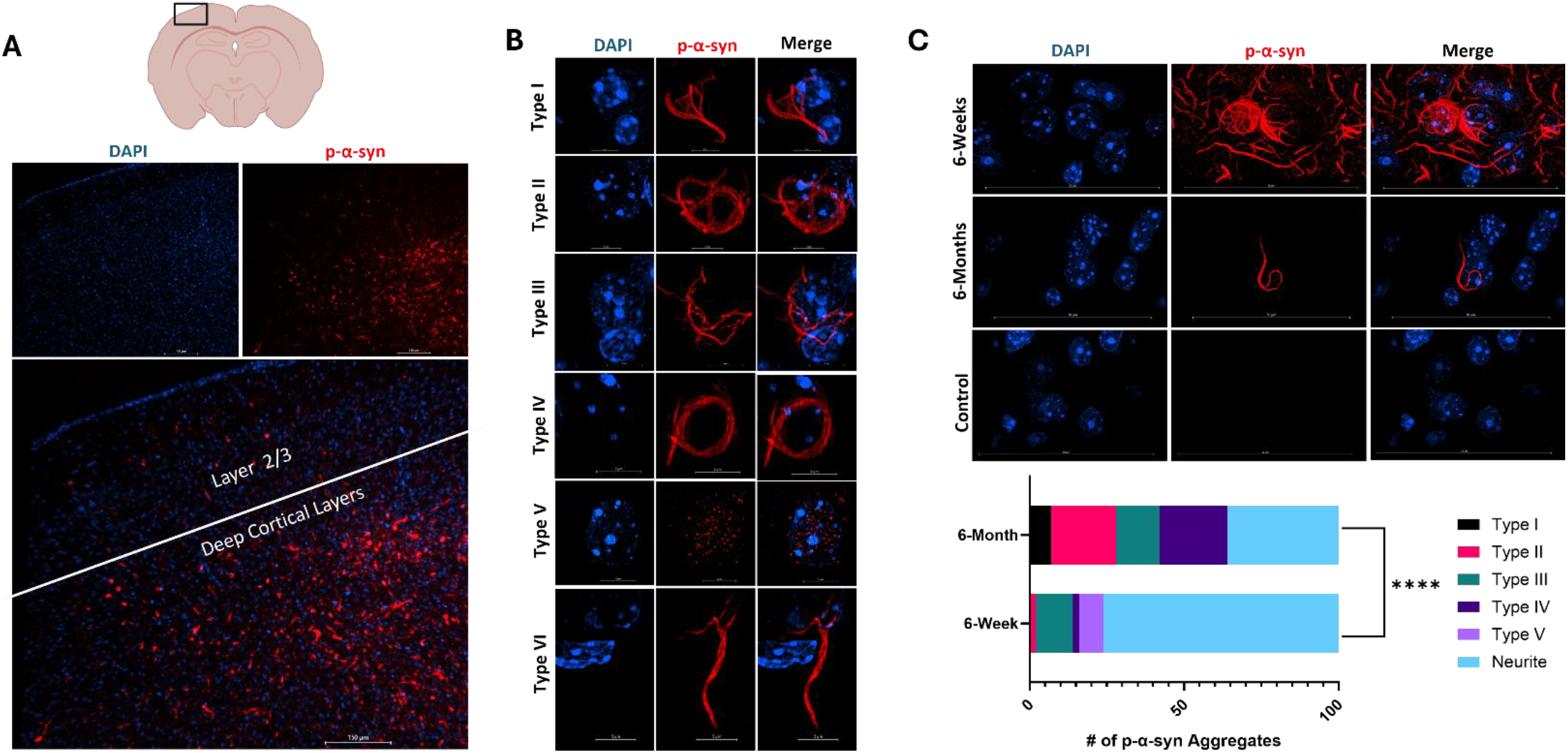
Synucleinopathy Mice Display Robust Cortical Lewy-like Pathology. A) Illustration depicting coronal slice of mouse brain tissue (Top), black box indicates example region of interest for imaging (Top). Representative micrograph of cortical Lewy-like Pathology at 10x magnification illustrating accumulation of phospho-Ser129-α-synuclein primarily in deep cortical layers (Bottom). B) Murine cortical Lewy-like pathology assumes six predominant morphologies. Ultrastructures of pathological α-synuclein took on 6 charecteristic morphologies.Type I ahhregaes a cell body filling structure that has pathological synuclein continuously extending into an axon or dendrite from the structure present in the cell body which often resembles a harp. Type II aggregates are a ring-like structure of pathological alpha-synuclein around the nucleus that has a loop connecting two sides of the ring structure into a “basket-like” structure. Type III aggregates are defined as multiple skein-like structures of alpha-synuclein that emanate from a common point that form a cage-like structure of alpha-synuclein in the cell body around the nucleus. Type IV aggregaes are defined as a ring-like structure of alpha-synuclein in the cell body. Type V aggregates are defined as >5 puncta within the nuclear compartment that do not form connected structures. Type VI aggregates are long rod like inclusions that are not present in the cell body but are present in dendrites and axons C) Pathological α-syn aggregation dynamics over time.

Examining the presence of nuclear interacting p-α-syn over time revealed a dynamic change in interaction with the nucleus. Using micrographs acquired both at 6-weeks and 6-months post PFF injection we found differential distribution of nuclear or perinuclear morphology. Using the criteria as established above, we tracked changes in morphology over time from 6 weeks post-injection to 6 months post-injection. These data show that p-α-syn aggregation is a dynamic process with significantly more punctate and neuritic pathology at 6 weeks and more fibril-like Type I through IV pathology at 6-months post PFF injection (p<0.0001, Fisher’s Exact Test) (Fig. 1C). There is an overall decrease in the amount of pathological α-syn structures per field of view in 6-month micrographs compared to 6-week micrographs (Fig. 1C), consistent with time-dependent pathology decreases seen in literature [50].Taken together, these data show that p-α-syn forms complex cell body ultra-structures in the peri-nuclear space which dynamically change over time.

Micrographs of representative pathology at 6-week and 6-month timepoints post PFF injection (Top). Quantification of aggregate morphology at 6-weeks and 6-months post PFF injection (Bottom). Of 100 aggregates analyzed at 6-weeks post PFF injection, we found; 0 Type 1, 2 Type II, 12 Type III, 2 Type IV, 8 Type V,and 76 Type VI(Neurites). Analysis of tissue harvested 6-months post PFF injection yielded; 7 Type 1, 21 Type II, 14 Type III, 22 Type IV, 0 Type V, and 36 Neurites. N= 3-6 mice, n=100 aggregates (p<0.0001, Fisher’s Exact Test).

### Pathological α-Syn Localizes Within Nuclear Compartments

Next, to determine if pathological aggregates of α-syn localize within the nucleus, we examined p-α-syn fluorescent signal localization within Lamin A/C (Abcam) structures for quantification. Lamin A/C was chosen as an internal “cutoff” structure to qualify if p-α-syn localizes within the nuclear compartments as it lines the inner most portion of the nuclear envelope (Fig. 2A, Left) [1]. Using IMARIS software (Oxford Instruments) we generated 3D surface renderings based off absolute fluorescent intensity to aid in quantification and visualization (Fig. 2B). This workflow was developed using mouse PFF injected tissue. Using this pipeline, we implemented a 240 nm lateral and 600 nm axial localization depth for p-α-syn within the nucleus as another “cutoff” metric for quantification to conservatively account for resolution limits of our image acquisition (Sup. Fig. 4). These cutoffs are both twice our established resolution limits, and establish conservative criteria to definitively call presence of p-α-syn inside the nuclear compartment or not. Utilizing these cutoff criteria, in mouse tissue, we observed multiple points where pathological α-syn could be seen passing through the nuclear lamina (green arrows), and colocalizing with DNA (black arrows) (Right magnified panels Figure 2B) Quantifying this across animals, we found 86.7% (95% CI 70.32-94.69%) of p-α-syn aggregates had substantial nuclear localization within the Lamin A/C structure (Fig. 2A, Right panel). See supplemental video 1 for animation of p-α-syn aggregation within nuclear compartments in mouse tissue.

**Figure 2:**
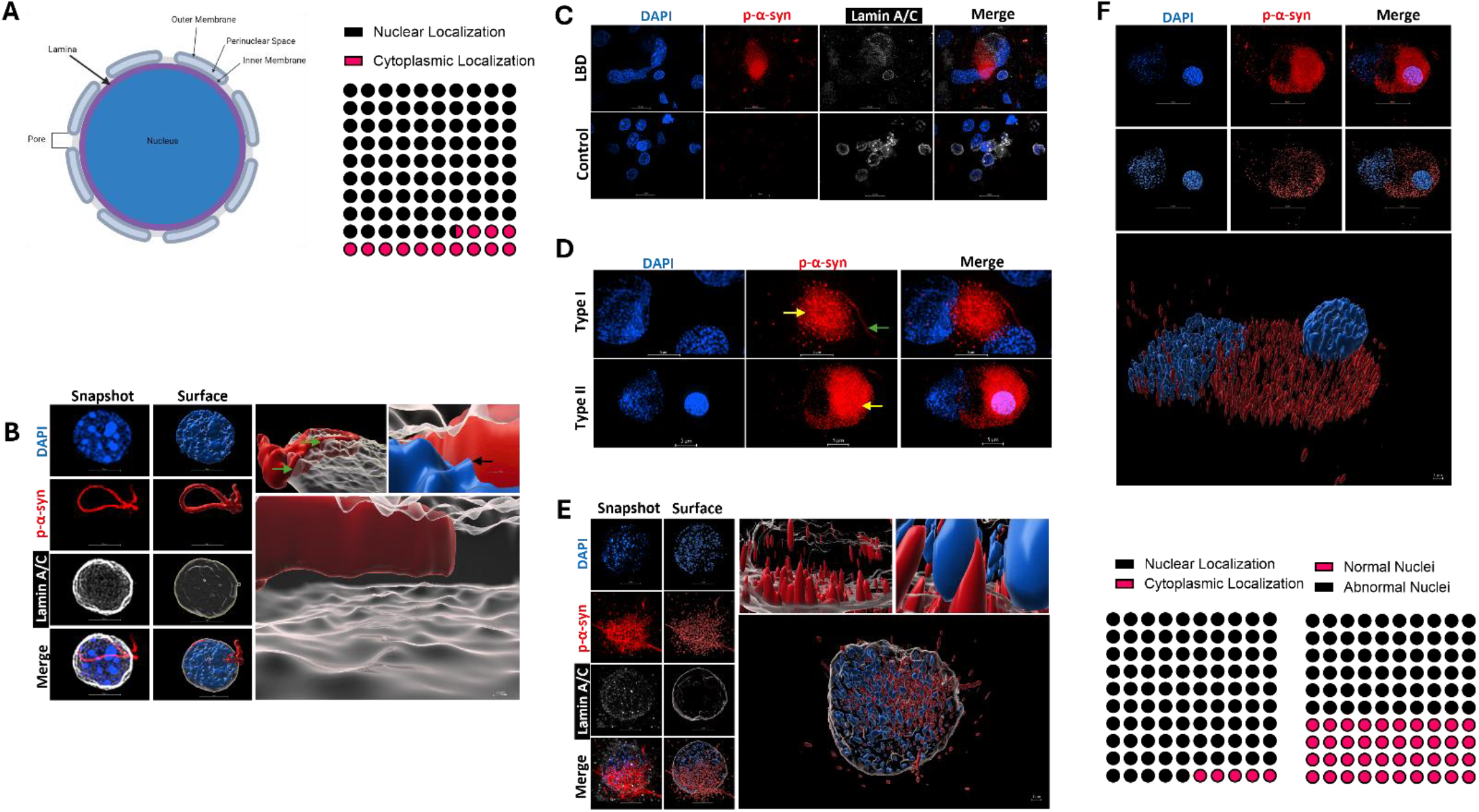
Pathological α-Syn Localizes Within Nuclear Compartments. **A)** Schematic depicting rudimentary nuclear envelope anatomy (Left). Nuclear Lamina (depicted in purple) proteins were used as a boundary structure to classify localization of α-syn within the nucleus, as it represents the innermost layer of the nuclear envelope. Quantification of p-α-syn localized withing the nuclear lamina yielded 86.7% (95% CI 70.32-94.69%) of aggregates with nuclear localization. N=3, n= 30 cells (p<0.0001, Binomial test). **B)** Images of DAPI, phospho-synuclein, and Lamin A/C were captured throughout the entire thickness of the cell body (Snapshot, left column, and were used to create surface renderings for each nuclear structure of interest and p-α-syn in IMARIS (Middle column)). These 3D surface renderings allowed for the visualization and quantification of p-α-syn interacting with the nuclear lamina(and an inside view of cell nucleus interacting with p-α-syn (Zoomed in surfaces, Top, Right). Green arrows indicate portions of the lamina where p-α-syn passes through the Lamin coating. Black arrow depicts colocalization of DAPI and p-α-syn. Bottom panel is a zoomed in view of inside the nuclear lamina which illustrates the continual presence of p-α-syn within this compartment. For a full animation of this panel sSee supplemental Video 1 for 3D animation of p-α-syn (Red) localizing within the nuclear lamina (White). (Animation of bottom right image). **C)** Representative 60x SoRa field of view micrographs of human cingulate cortex tissue. Lewy Body Dementia (LBD) tissue (Top) and control tissue (Bottom) using the same staining paradigm as in (B). **D)** Lewy Pathology observed were punctate aggregates (Yellow arrow)either with (Type I) (Top) or without (Type II) (Bottom) an attached fibril-like component (Green arrow). **E)** Lamin surfaces were generated for visualization purposes on high integrity nuclei as we did in (B), zoomed in surfaces revealing rod-like and punctate p-α-syn structures throughout the nucleus (Right panels). Bottom panel is a still shot of See supplemental Video 2A which shows a 3D animation of p-α-syn (Red) localization within an intact lamina (White). **F)** Nuclear localization was quantified by co-localization of p-α-syn with DAPI, shown in representative Type II Lewy Body from Fig. 2D (Bottom) Quantification of nuclear localization (Bottom, Left) revealed 95% (95% CI76.39 to 99.74) of aggregates analyzed have nuclear localization (p<0.0001, Binomial test). N= 3 LBD Cingulate cortices, n= 20 aggregates. Of the nuclei these aggregates interacted with, 60% (95% CI 38.66 to 78.12) displayed aberrant morphology (Bottom, Right) (p<0.0001, Binomial test). N= 3 LBD Cingulate cortices, n= 20 aggregates. See supplemental Video 3 for 3D animation of p-α-syn (Red) pathology interacting with DAPI (Blue).

To determine if pathological p-α-syn is present in the nucleus in human synucleinopathies, as previously suggested, [25], we processed and analyzed control and Lewy Body Dementia (LBD) tissue as described for murine tissue with minor modifications (see Materials and Methods). See table 2 for detailed information on human tissue employed in this study. Human LBD Lewy Pathology was primarily neuritic with high density of small punctate, shapes, and neuritic morphology (Fig. 2C, Sup. Fig. 5). We do not observe any appreciable accumulation of pathological p-α-syn in the cell bodies of human donors without synucleinopathies. The large cortical Lewy pathology analyzed were predominantly accumulations of small p-α-syn aggregates (yellow arrows in figure), but some have fibril-like components attached to the punctate mass (green arrows, Fig. 2D). Cortical pathology in patients with synucleinopathies fell into two categories. Those with accumulations of small aggregates and had fibril like components (Type I), and those that exclusively had accumulations of small aggregates (Type II) Both control and LBD tissue had some level of Lamina damage and/or degradation (Fig. 2C, Sup. Fig 3) that could likely be attributed to the postmortem interval (PMI) before tissue fixation (See Table 2 for PMI data) [5, 7]. Due to varied levels of nuclei integrity (Sup. Fig. 3), Lamin surface renderings were only generated for visualization purposes on high integrity nuclei (Fig. 2E). See supplemental video 2 for animation. Cortical Lewy pathology is perinuclear with significant levels of nuclear localization as well in patients with synucleinopathies (Fig. 2C-F). Analyzing cortical Lewy pathology for co-localization with DAPI in our patient panel, we found nuclear localization to also be a hallmark of LBD, with 95% of these aggregates having nuclear localization (Fig. 2F). When examining if this nuclear localization of the aggregates alters neuronal morphology, we found that 60% (95% CI 38.66 to 78.12) of nuclei that contained pathological p-α-syn were associated with nuclear abnormalities (Fig. 2F, bottom right). These data suggest that Lewy pathology is present in human synucleinopathies as well, and contributes to envelope irregularities. These data further suggest that the nuclear localization of pathological p-α-syn in the rodent PFF model is not an artifact of that model, and that nuclear localization of pathological p-α-syn in cortical neurons may be an important phenotype of synucleinopathies. See supplemental video 3 for animation of Lewy pathology colocalization with DNA in patients with synucleinopathies.

**Table 2:** Demographic Information and Neuropathological Diagnoses of Patients with Synucleinopathies: Age, Gender, Post-Mortem Interval, and Neuropathological characterization of human donor tissue used in this study.

|  | Neuropathological Diagnosis | Age (years) | Gender | PMI (hours) | Thal Phase | Braak | CERAD |
| --- | --- | --- | --- | --- | --- | --- | --- |
| Control |  |  |  |  |  |  |  |
|  | Cerebrovascular Disease | 73 | M | 48 | 1 | I | None |
|  | No Significant Findings | 53 | M | 72 | 0 | 0 | None |
| | Diffuse Aβ Pathology | 65 | F | 15 | 1 | I | None |
| Synucleinopathy |  |  |  |  |  |  |  |
|  | Dementia with Lewy Bodies | 79 | M | 12 | 5 | VI | Frequent |
|  | Parkinson's Disease | 63 | M | 27 | 1 | II | None |
|  | Parkinson's Disease Dementia | 82 | M | 12 | 3 | II | Sparse |

### Nuclear p-α-Syn Induces Nuclear Abnormalities

Because of the limiting factor of Lamin A/C and nuclear integrity damage being related to PMI and not necessarily pathological p-α-syn withing the nucleus (Sup. Fig 3), we rigorously characterized the effects of p-α-syn nuclear localization on nuclear integrity using our rodent model. This rodent displays similar nuclear localization of pathological p-α-syn to patients. We observe multiple nuclear envelope irregularities in cortical mouse synucleinopathy tissue which we found to have 3 primary phenotypes. These are classified as multi-lobed (multiple lamina coated structures encased by a single Lewy aggregate), malformed (altered Lamina structure as compared to control that is still a contiguous structure), or ruptured (nuclear Lamina has a “hole”, or is no longer a contiguous structure, Fig. 3A). Nuclei were frequently found to have more of these phenotypes but were classified as the more predominant irregularity in our analysis. Qualitative assessment of atypical phenotypes observed, showed they occur at roughly equal frequency to one another, with 31% (95% CI 11.79-40.93) multi-lobed, 30% (95% CI 9.51-37.31) malformed, and 35% ruptured (Fig. 3B, Left).This is similar to our findings in human tissue that indicate 60% of cortical nuclei with pathological p-α-syn show irregular nuclei (Figure 2 C-F).

**Figure 3:**
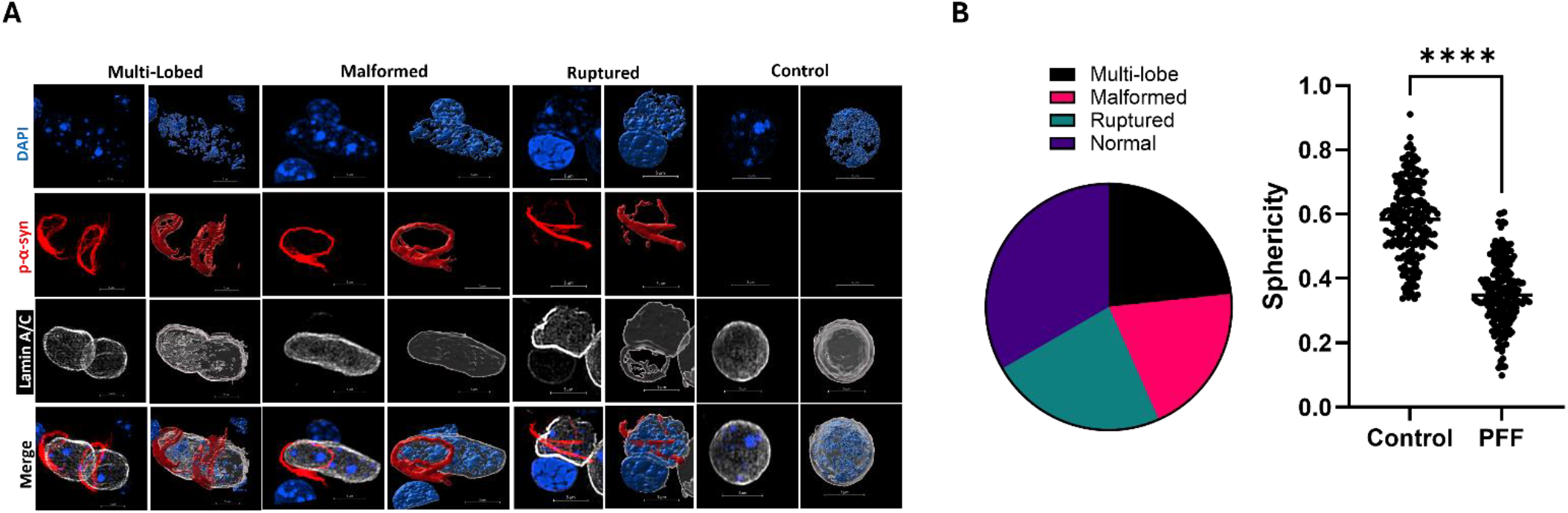
Nuclear p-α-Syn Induces Nuclear Abnormalities. **A)** Example micrographs of mouse nuclei from control or synucleinopathy neurons that exhibit abnormal nuclei morphologies. Abnormalities broadly fell into three categories; multi-lobed, malformed, and/or ruptured. Fluorescent signal is shown on the left and surface rendering on right for each classification. Representative control nuclei shown (Right Column). N= 3-6 mice, n=50 nuclei. **B)** Distribution of abnormal nuclei classifications in cortical synucleinopathy tissue (Right) from the cells examined in Fig. 3A. Of the abnormal nuclei; 35% (95% CI 11.79-40.93) of nuclei were multi-lobed, 30% (95% CI 9.51-37.31) were malformed, and 35% (95% CI 11.79-40.93) appeared ruptured (Right). Sphericity(ψ) of cortical cell quantified in control and synucleinopathy tissue (Left). Control tissue averaged ψ=0.5836 (95% CI 0.5678-0.5994) and PFF treated tissue averaged ψ=0.3550 (95% CI 0.3414-0.3687) N= 3-6 mice, n= 210 cells (p <0.0001, Unpaired-T test, t=21.60).

To quantitatively assess nuclear integrity, we next quantified sphericity of nuclear lamina structures. Sphericity (ψ) is the measure of how close a given object is to a true sphere with a maximum value of 1 [58]. We quantified the sphericity in synucleinopathy and control murine tissue which yielded an average of ψ=0.3550 (95% CI 0.3414-0.3687) and ψ=0.5836 (95% CI 0.5678-0.5994) respectively (Fig. 3B, Right). Nuclei with p-α-syn show a significant decrease in nuclear sphericity (p<0.0001, Unpaired-T test, t=21.60). This suggests loss of nuclear integrity in synucleinopathy tissue can be measured both qualitatively using our established IMARIS imaging pipeline, as well qualitatively.

### Synucleinopathy Models have Increased Genotoxicity

We predicted loss of nuclear integrity and altered nuclear morphology in cortical synucleinopathy tissue was associated with increased genotoxicity as there are reports of increase DNA damage and genotoxicity in synucleinopathy models of other brain regions [35, 56]. Murine tissue was probed with genotoxic and DNA damage marker tumor suppressor P53-binding protein 1 (53BP1). 53BP1is associated with physical squeezing and damage to the nuclear envelope [22, 53], which may result from p-α-syn aggregate interaction with the nucleus. An increase in nuclear 53BP1 puncta was seen throughout the cortex, both in cells with and without α-syn aggregation (Fig. 4A left). Due to species mismatch, we utilized a total-synuclein antibody for Figure 4A, but cells with aggregates were easily identifiable still due the morphology of α-synuclein. Quantitation of nuclear 53BP1 levels revealed significantly more (p<0.0001, Unpaired-T test, t=5.215) 53BP1 puncta in mouse synucleinopathy tissue, with an average of 66.89 puncta per cell (95% CI 54.89-78.89) while control cortical cells averaged 29.64 (95% CI 22.10-37.18) 53BP1 puncta per cell (Fig. 4A, Right).

**Figure 4:**
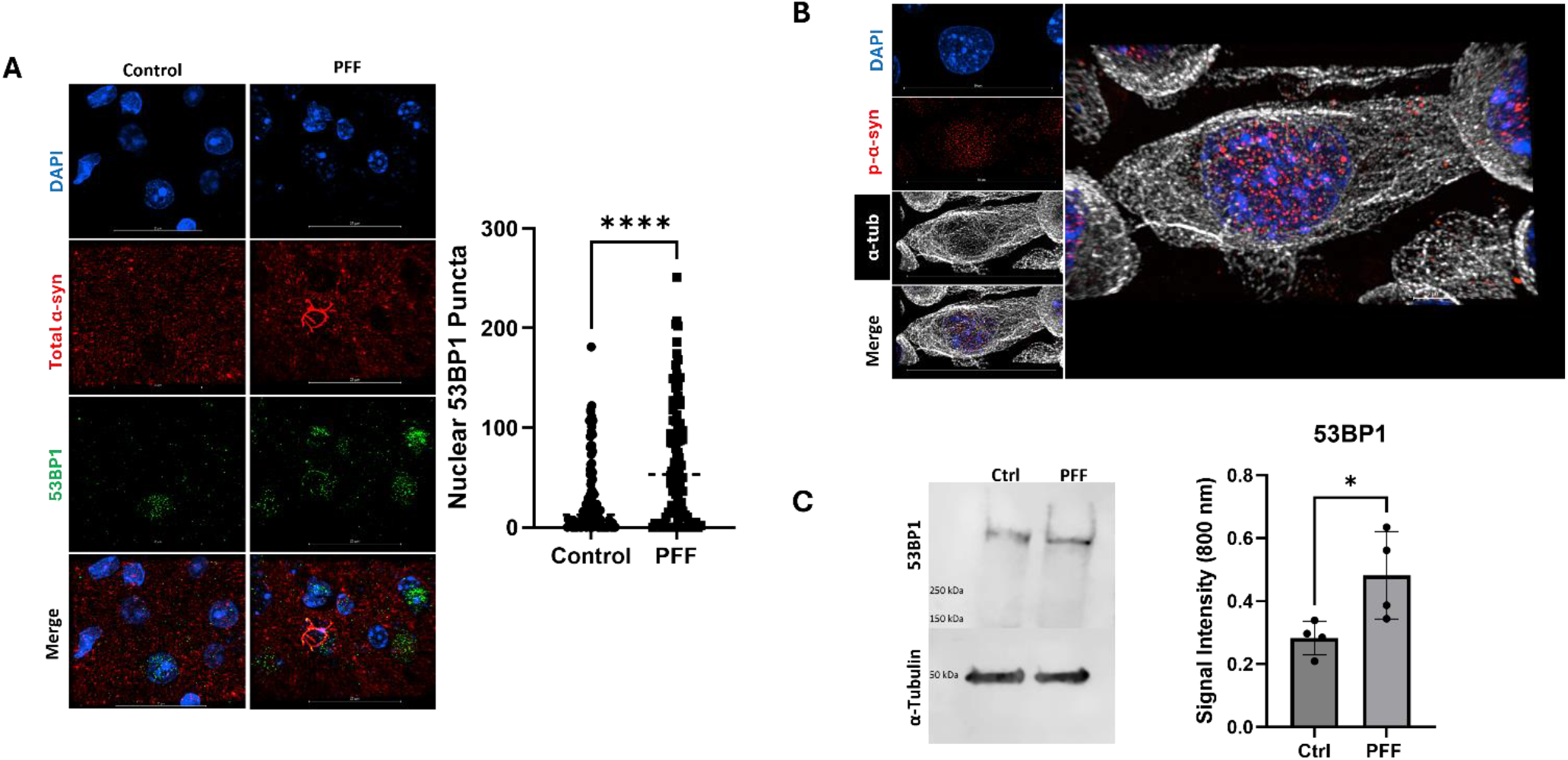
Synucleinopathy Models Have Increased Genotoxicity. **A)** Murine cortical neurons display increased 53BP1 puncta compared to wild type tissue. Quantification (Right) yielded an average of 66.89 puncta (95% CI 54.89-78.89) per synucleinopathy cell while control cortical cells averaged 29.64 53BP1 puncta per cell (95% CI 22.10-37.18).N=3-6 mice, n=100 cells (p<0.0001, Unpaired-T test,t=5.215). **B)** Neuro 2a cells treated with 5.72 µM PFFs via the BioPORTER protein delivery system, accumulate pathological α-syn in nucleus with diffuse p-α-syn also observed throughout the cell. See supplemental Video 4 for 3D animation of p-α-syn (Red) inclusions localizing throughout the cell body (White, α-tubulin) of Neuro 2a cells but with highest inclusion concentration within the nucleus (DAPI, blue). **C)** Nuclear accumulation of p-α-syn in Neuro 2a cells is associated with increased 53BP1 genotoxicity marker, with signal intensity ratio of 53BP1 dived by α-tubulin averaging 0.2826 (95% CI 0.1971-0.3682) and 0.4821 (95% CI 0.2612-0.7029) in control (Ctrl) and PFF treated cells respectively… N=4 biological replicates (p=0.0366, Unpaired-T test, t=2.68).

Next, we employed a cellular model of nuclear p-α-syn accumulation in synucleinopathies to define the downstream effects of nuclear localization on the cell nucleus. Adapting upon previous cell models [51], we used protein transfection of PFFs in murine Neuro 2a cells, which resulted in the robust accumulation of small p-α-syn punctate aggregates in the nucleus of the cell (Fig. 4B,Sup. Fig. 6), resembling Type V pathology in murine tissue (Fig. 1B) and human Lewy Pathology (Fig. 2C). See supplemental video 4 for animation of Neuro 2a nuclei with pathology. Nearly all Neuro 2a cells in culture display nuclear pathology (Sup. Fig. 6), enabling population assays. Consistent with our immunofluorescence results, western blot analysis of Neuro 2a cells for 53BP1 showed control cells display an average 53BP1 fluorescent signal ratio of 0.2826 (95% CI 0.1971-0.3682), while PFF treated cells averaged 0.4821 fluorescent signal ratio for 53BP1 signal (95% CI 0.2612-0.7029) (Fig. 4B, Bottom). Signal differences were significant (p=0.0366, Unpaired-T test, t=2.68).

### Pathological α-Syn Accumulation Results in Nuclear Deficits

To determine the functional impact of p-α-syn nuclear accumulation on the nucleus, we examined multiple measures of nuclear function using the same cell culture model as above in Figure 4B, First, we probed RNA localization, as impaired nucleo-cytoplasmic export has been suggested in synucleinopathies and other related neurodegenerative disorders [10, 60]. Analysis of RNA volume renderings, from labeling with a commercially available RNA specific dye (see table 1), showed a nuclear to cytoplasmic ratio of 0.4954 (95% CI 0.3508-0.5679) for synucleinopathy cells, and 0.09499 (95% CI0.02112-0.1689) for control tissue, demonstrating a significant decrease of RNA outside the nuclei of synucleinopathy cells (Fig. 5A) (p=0.0003, Unpaired-T test, t =11.94). To ensure that these changes in RNA localization are not due to changes in cellular morphology induced by PFF addition, we examined total cellular volumes using an α-tubulin stain which allowed us to create a complete volume rendering of each cell in IMARIS. We found that there are no differences in cell volume between PFF treated cells and control (Fig. 5B) (p=0.5397, Unpaired T-test, t=0.6697). Controls cells displayed an average α-tubulin staining volume of 1173 µM^3^ (95% CI 922.8-1424), while PFF treated cells averaged 1274 µM^3^ of α-tubulin volume per cell (95% CI 677.1-1871) (Fig. 5B). This data suggests nuclear p-α-syn negatively impacts RNA export, but further studies will be necessary to determine the mechanism that results in abnormal RNA localization.

**Figure 5:**
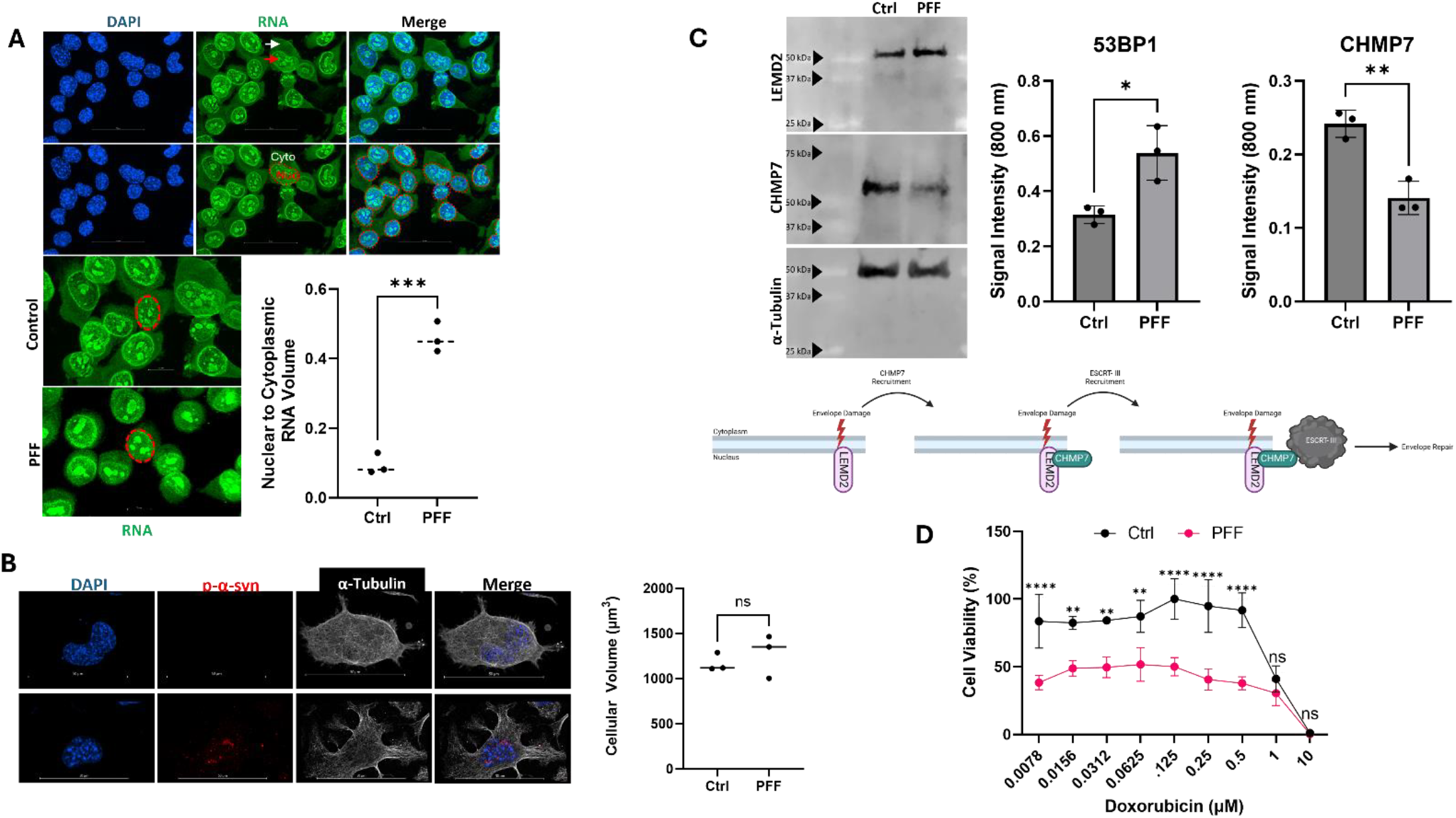
Pathological α-Syn Accumulation Results in Nuclear Deficits. **A)** RNASelect dye labeling of Neuro 2a cells clearly defines nucleoli (Red Arrow) and nuclear borders (Red Dotted Line). All fluorescent signal outside of the nuclear border was considered cytoplasmic (White Arrow). Quantification of staining volume distribution between nucleus and cytoplasm (Bottom) in control (Ctrl) and PFF treated cells yielded an of 0.4954 (95% CI 0.3508-0.5679) for PFF treated cells, and 0.09499 (95% CI0.02112-0.1689) for control treated cells. N=3 biological replicates, n >100 cells per replicate (p=0.0003, Unpaired-T test, t =11.94). B) Micrograph of immunofluorescence staining for α-tubulin in cells with (Bottom Row) and without (Top Row) p-α-syn pathology. Quantification of α-tubulin volume per field of view, normalized to number of nuclei per field of view, revealed an average cell volume of 1173 µm^3^ (95% CI 922.8-1424) for control cells and 1274 µm^3^(95% CI 677.1-1871) for PFF treated cells. N= 3 biological replicates (p=0.5397, Unpaired-T test, t= 0.6697). C) Western blot analysis of envelope repair pathway proteins in control and PFF treated Neuro 2aA cells. Alpha-tubulin (α-tubulin)(DM1A Abcam) loading control shown on bottom. Quantification of western blots (Right panel). Control treated cells displayed an average of 0.05319 signal intensity ratio for LEMD2 (95% CI 0.02738-0.07901), while PFF treated cells averaged 0.1080 fluorescent signal ratio for LEMD2 (95% CI 0.05362-0.1624) (p=0.0173, Unpaired-T test, t=3.918). Control cells yielded an average CHMP7 (Proteintech) signal intensity ratio of 0.2419 (95% CI 0.1961-0.2877) compared to 0.1409 signal intensity ratio (95% CI 0.08469-0.1971) for CHMP7 in PFF treated cells(p=0.0039, Unpaired-T test, t=5.994). Visual rendering of simplified ESCRT-III mediated envelope repair pathway (Bottom). **D)** Neuro 2aA cells were treated with increasing concentrations of the DNA damaging agent, Doxorubicin, for 24 hours. Cell viability was determined with Cell Titer Glo 2.0 assay and normalized to untreated wells, and expressed at percent of the well with the highest luminescent signal. No statistical differences (Two-way ANOVA) were observed at 10 µM doxorubicin (1.10% Ctrl, 0.35% PFF Viability), or at 1 µM doxorubicin (41.12% Ctrl, 30.43% PFF viability). Statistical differences (Two-way ANOVA) in viability between control and PFF treated cells after doxorubicin treatment were present at 0.5 µM doxorubicin (91.64% Ctrl, 37.84% PFF viability) (Mean Diff.= 0.6929, 95% CI of Diff. 0.3744-1.011, p<0.0001), 0.25 µM doxorubicin treatment (94.79% Ctrl and 40.51% PFF viability) (Mean Diff= 0.6991, 95% CI of Diff. 0.3806-1.018, p<0.0001), 0.125 µM doxorubicin (100% Ctrl, 50.04% PFF viability) (Mean Diff.= 0.6439, 95% CI of Diff. 0.3254-0.9625, p<0.0001), 0.0625 µM (87.15% Ctrl, 51.71% PFF viability) (Mean Diff.=0.4565, 95% CI of Diff. 0.1379-0.7750, p=0.0024), 0.0312 µM doxorubicin (84.24% Ctrl, 49.60% PFF viability) (Mean Diff.= 0.4462, 95% CI of Diff. 0.1276-0.7647, p= 0.0019), 0.0156 µM (82.45% Ctrl, 48.85% PFF viability) (Mean Diff.= 0.4328, 95% CI of Diff.0.1143-0.7514, p= 0.0027), and 0.0078 µM (83.62% Ctrl, 38.31% PFF viability) (Mean Diff.=0.5836, 95% CI of Diff. 0.2650-0.9021, p<0.0001). We previously determined that PFF treatment alone does not decrease viability of Neuro 2A cells.). N=3 biological replicates, n= 6 technical replicates. See table 1 for a complete list of all antibodies and reagents.

Second, we investigated the impact of nuclear p-α-syn on envelope repair machinery, namely endosomal sorting complexes required for transport (ESCRT-III) repair pathway proteins (Fig. 5C). Western blots were performed on synucleinopathy and control Neuro 2a cells reveals differential expression of LEMD2 and the ESCRT-III component CHMP7 [19] (Fig. 5C). LEMD2, which accumulates at nuclear envelope damage/ruptures and recruits CHMP7 to initiate repair [19], was found to have an average fluorescent signal ratio of 0.05319 in control cells (95% CI 0.02738-0.07901), while PFF treated cells averaged 0.1080 signal intensity ratio for LEMD2 signal (95% CI 0.05362-0.1624) (p=0.0173, Unpaired-T test, t=3.918) Moreover, Control cells yielded an average CHMP7 signal intensity ratio of 0.2419 (95% CI 0.1961-0.2877) compared to a fluorescent signal ratio of 0.1409 (95% CI 0.08469-0.1971) in PFF treated cells (p=0.0039, Unpaired-T test, t=5.994). This suggests that there is increased nuclear envelope damage, but a diminished or dysfunctional capacity to attempt to repair this damage.

### Nuclear p-α-syn Sensitizes Cells to Nuclear Toxin Insult

Lasty, we examined whether decreased nuclear integrity affected susceptibility to nuclear insult and subsequent cell death. Control or PFF inoculated Neuro 2a cells were treated with increasing concentrations of doxorubicin, a DNA intercalator that inhibits topoisomerase II [3, 14, 47], for 24 hours and cell viability was subsequently recorded using a Cell Titer Glo Assay (see table 1). Viability after 24 hours of doxorubicin treatment at concentrations less than 0.5 μM were significantly lower in cells with nuclear pathological α-syn (Fig. 5D) (Two-way ANOVA). At 0.5 µM doxorubicin control cells averaged 91.64% viability and PFF treated cells 37.84% viability (Mean Diff.= 0.6929, 95% CI of Diff. 0.3744-1.011, p<0.0001). After 0.25 µM doxorubicin treatment, control cells averaged 94.79% viability and PFF treated cells averaged 40.51% viability (Mean Diff= 0.6991, 95% CI of Diff. 0.3806-1.018, p<0.0001). At 0.125 µM doxorubicin, control cells averaged 100% viability, while PFF treated cells yielded 50.04% viability (Mean Diff.= 0.6439, 95% CI of Diff. 0.3254-0.9625, p<0.0001). Doxorubicin treatment at 0.0625 µM resulted in a control cell mean viability of 87.15% compared to 51.71% for PFF treated cells (Mean Diff.=0.4565, 95% CI of Diff. 0.1379-0.7750, p=0.0024). Viability after 0.0312 µM doxorubicin treatment was 84.24% for control cells and 49.60% in PFF treated cells (Mean Diff.= 0.4462, 95% CI of Diff. 0.1276-0.7647, p= 0.0019). Doxorubicin treatment at 0.0156 µM resulted in 82.45% average control viability and 48.85% average viability of PFF treated cells (Mean Diff.= 0.4328, 95% CI of Diff.0.1143-0.7514, p= 0.0027). The lowest concentration of doxorubicin tested was 0.0078 µM, which resulted in average control viability of 83.62% and 38.31% (Mean Diff.=0.5836, 95% CI of Diff. 0.2650-0.9021, p<0.0001). This suggests the presence of nuclear α-syn sensitizes the cell to death due to environmental insults.

## Discussion

The nuclear localization of α-syn has remained controversial despite numerous reports of α-syn in the nuclei of various animal and cellular models both in pathogenic states and normal physiology (Goers et al., 2003; Liu et al., 2011; Maroteaux et al., 1988; McLean et al., 2000; Mori et al., 2002; Pinho et al., 2019; Schaser et al., 2019; Yu et al., 2007). Lack of nuclear characterization and off target nuclear staining associated with early α-syn antibodies may have contributed to this controversy [20, 25]. Studies have also forced nuclear localization of α-syn via adding a nuclear localization sequence and observed downstream detrimental nuclear function due to increased nuclear α-syn [12, 46], suggesting α-syn is a critical regulator of nuclear function. Recently, pathological α-syn has been suggested in postmortem LBD tissue [25], raising questions as to potential pathological nuclear function and the downstream impact of nuclear p-α-syn. Using quantitative super resolution imaging, we rigorously report robust nuclear p-α-syn aggregates in our synucleinopathy mouse model, cell culture model, and postmortem human tissue. This nuclear localization is associated with functional and morphological nuclear abnormalities that raise a number of implications for future therapeutic avenues as well as pathogenesis of disease. Taken together, these data have led us to a working model (Figure 6) in which presence of pathological α-syn in the cell body directly interacts with the nucleus, impairs nuclear morphology, alters neuronal function, and these changes sensitize the neuron to environmental insults leading to cell death.

**Figure 6:**
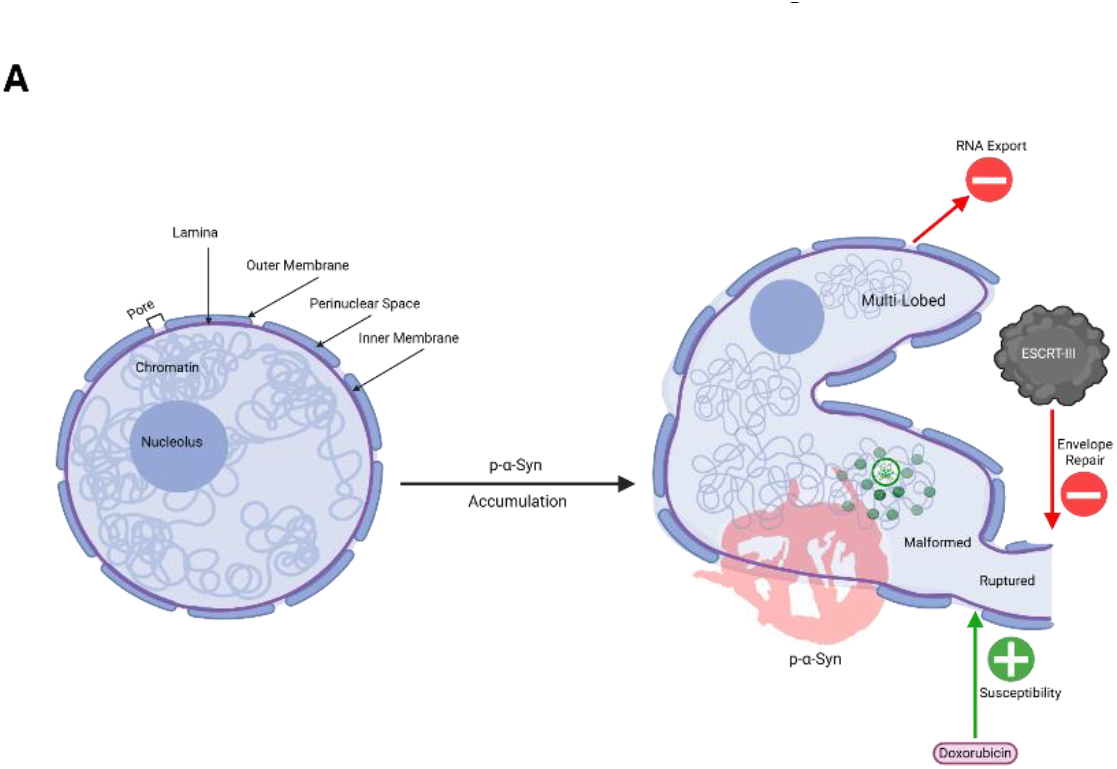
Intranuclear p-α-Syn Compromises Integrity and Function of the Nucleus. **A)** Visual rendering of our working hypothesis on pathological α-syn nuclear localization and downstream consequences of this localization. The nucleus on the left depicts a healthy neuronal nucleus without pathological α-syn. The major components of the nucleus are labeled. Upon nuclear accumulation of p-α-syn, abnormal nuclear morphology is apparent. Namely rupturing (depicted as discontinuation in membrane and lamina), malformation (depicted as overall decrease in sphericity of nuclear morphology), and formation of multiple lobes (depicted as smaller lobe branching off the main nuclear lobe). Abnormal morphology is correlated with an increase in 53BP1 genotoxicity (Green Puncta), decreased ESCRT-III component expression (which we predict decreases envelope repair as a result)(Middle Red Arrow), aberrant RNA localization (which we predict is due to impaired RNA export) (Top Red Arrow), and increased sensitivity to environmental insult, specifically doxorubicin (Bottom Green Arrow).

The abnormal nuclear morphological phenotypes we observed in synucleinopathy tissue from human and mouse closely resembles the morphological abnormalities observed in laminopathies, a group of disorders resulting from mutations in the *LMNA* and *LMNB* genes that encode the nuclear lamina proteins Lamin A/C and Lamin B respectively [8, 61]. Laminopathies are associated with abnormal nuclei morphology and loss of nuclear integrity [49, 61]. We observed increased abnormal nuclei morphology in both murine and human synucleinopathy tissue, suggesting a common phenotype between synucleinopathies and laminopathies (Fig. 2F, 3).Consistent with reports of α-syn induced nuclear damage in other synucleinopathy models and brain regions [35, 56], we report for the first time, to our knowledge, that cortical neurons *in vivo* and cultured Neuro 2a cells post α-syn corruption by PFFs have increased 53BP1 nuclear localization (Fig. 4A). 53BP1 is associated with a specific form of DNA damage that is closely linked with physical stress placed on the nucleus [22, 53], correlating morphological changes in nuclear morphology to genotoxicity. Laminopathies are also associated with increased DNA damage and genotoxic markers [13, 18], drawing further similarity between laminopathies and synucleinopathies described in our models. Our data directly suggests that nuclear localization of pathological α-syn causes physical damage to the nuclear envelope, resulting in morphological changes associated with impaired function.

Our data further shows impaired nuclear function as a result of the nuclear damage described above. First, we observed apparent functional deficits in our cell culture model by labeling cells with an RNA specific dye. Synucleinopathy cells were found to contain significantly less cytoplasmic volume of RNA staining compared to controls (Fig. 5A). Impaired mRNA export similar to that we have observed has been found to result from aggregation or dysfunction TAR DNA binding protein-43 (TDP-43) in other neurodegenerative disorders, namely ALS. Additionally, neuronal aggregation of TDP-43 and subsequent mRNA dysfunction is hypothesized to occur in other neuronal protein-aggregation disorders, including genetic forms of PD [60]. Impaired protein export [10] has also been reported in pathogenic SCNA models. Our RNA labeling experiment suggests that RNA export is impaired in cells with α-syn pathology due to lack of cytoplasmic staining, suggesting impaired RNA transport out of the nucleus (Fig. 5a, 6). Future studies will aim to elucidate the mechanistic impact of nuclear p-a-syn pathology on *Nxf1* dependent export of mRNA [57], Exportin-t dependent export of tRNA [26] and CRM1-dependent rRNA export [6].

Additional nuclear dysfunction was implicated via analysis of nuclear envelope repair pathways, that we investigated due to initial findings of envelope disfigurement and ruptures in nuclear lamina (Fig. 3A). The endosomal sorting complex required for transport III complex (ESCRT-III) is well characterized in envelope repair [48, 59]. Within the envelope repair pathway (Fig. 5C), LEMD2 localizes to envelope damage and recruits ESCRT-III component CHMP7, that subsequently recruits other ESCRT-III proteins [19]. We interpret the observed increased LEMD2 expression but decreased CHMP7 expression as a likely increase in nuclear envelope damage but impaired ESCRT-III mediated repair. Future studies will aim to fully characterize the impact of nuclear p-α-syn on envelope repair mechanisms, how this contributes to our observed deficits, and whether targeting the repair pathway therapeutically could mitigate changes in nuclear function.

Finally, we investigated the impact of p-α-syn induced nuclear deficits on cell survival. Both in cell culture and rodent synucleinopathy models, accumulation of p-α-syn into aggregates was not directly associated with decreased viability. Therefore, we examined whether p-α-syn sensitizes cells to environment toxin induced cell death. Utilizing the nuclear damaging reagent doxorubicin, which alone has previously been shown to increase α-syn aggregation [16], we showed that cells with p-α-syn demonstrated decreased viability compared to control cells in response to low doxorubicin concentrations (Fig. 5D) [21]. This is of interest, as increasing reports of environmental factors, such as pesticide exposure, increase synuclein aggregation and risk for developing synucleinopathies [31, 40, 41]. This may give further support for a “two-hit” component in which genetic predisposition to α-syn aggregation, or accumulation of p-α-syn for other reasons, in combination with environmental stressors leads to or exacerbates diseases [55]. We predict that nuclear damage induced by p-α-syn aggregation sensitizes neurons bearing aggregates to environmental insults, and that targeting nuclear based dysfunction caused by p-α-syn may be a novel mechanism by which to prevent cell death in synucleinopathies.

Cumulatively, our data demonstrates the presence of nuclear pathological α-syn at super resolution detail and reveals atypical downstream nuclear phenotypes resulting from this disease associated localization. The prevalence of the nuclear p-α-syn phenotype, observed impact on nuclear integrity, and functional impairment of the nucleus, altogether suggest a relatively unexplored role for p-α-syn to disease etiology and cellular dysfunction in synucleinopathies. Additional mechanistic characterization of how p-α-syn localizes to the nucleus is necessary, and the full repercussions of nuclear p-α-syn are just beginning to be elucidated. Lastly, as different neuroanatomical regions respond uniquely to nuclear insults [2, 11, 29, 38], we predict α-syn induced nuclear stress will impact various brain regions differently, and understanding disease signaling within and between brain regions may provide mechanistic insight into synucleinopathies and cell type selective vulnerability.

## Conclusion

This study provides super resolution evidence of Lewy body and Lewy-like pathology localizing within the nuclei of non-pretreated human and murine tissue. We show nuclear localization is correlated to increases in abnormal nuclear envelope morphologies. Additionally, a cellular model of nuclear pathological α-syn displayed abnormal RNA localization and differential expression of envelope repair pathway proteins, suggesting altered functionality of the nucleus in cells with pathology. Lastly, we show cells with nuclear Lewy-like pathology are more susceptible to nuclear insults, suggesting a potential two-hit component in which environmental factors influence disease etiology.

## List of Abbreviations

α-syn: Alpha-synuclein
p-α-syn: Pathological alpha-synuclein
LBD: Lewy body dementia
PFF: Pre-formed fibril.
PBS: Phosphate buffered saline
DNA: Deoxyribonucleic acid
RNA: Ribonucleic acid
LEMD2: LEM domain nuclear envelope protein 2
CHMP7: Charged multivesicular body protein 7
ESCRT-III: Endosomal sorting complexes required for transport-III
SEC: Size exclusion chromatography
AEC: Anion exchange chromatography
EDTA: Ethylenediaminetetraacetic acid
ANOVA: Analysis of variance
Ctrl: Control
TDP-43: TAR DNA binding protein-43

## Declarations

### Ethics

All animal work in this manuscript was approved by the University of Florida Institutional Animal Care and Use Committee before starting the work. Human tissue work was approved by an exemption from the University of Florida Institutional Review Board.

### Data Availability

All data from this paper are able to be requested from the corresponding author upon reasonable request.

### Author Contributions

MM, AC, EMC, AH, PW, IG, JR, NC DH, EM, DN, and BB designed experiments, performed experiments, collected data, and analyzed data. SP provided human tissue samples and associated pathological characterization. SP and MSM provided funding. MM wrote the first draft of the manuscript. MM, SP, and MSM edited the manuscript to the final version. MSM and SP provided supervision of the manuscript. All authors read the manuscript, provided feedback, and approved the final version.

## Acknowledgments

We would like to thank the laboratory of Dr. Jonathan Bird for providing access and technical assistance with Nikon CSU-W1 Spinning Disc with Gataca super resolution modules. We would also like to thank Bird laboratory member James Heiding’s, for expert liquid chromatography technical support. We additionally acknowledge the UF Center for Translational Research in Neurodegenerative Disease (CTRND) for access to and maintenance of Nikon SORA super resolution microscope and cell culture facilities, and to Dr. Michael Goedert for gifting us the PRK172-alpha-synuclein plasmid. We sincerely thank the patients and families for the generous donation of human tissue used in this study.

## Supplemental Figure Legends

**Supplemental Figure 1: Quality Assurance Metrics for α-Synuclein Purification from BLD21 cells expressing PRK172 plasmid. A)** Gelcode Blue stained SDS-PAGE gels loaded with probable fractions generated from liquid chromatography. Top image depicts fractions after anion exchange, with nine total positive for α-syn.. Bottom image depicts probable fractions after size exclusion chromatography, with four fractions positive for α-syn (fractions 4-7). **B)** Representative dynamic light scattering (DLS) sizing quality check of PFF samples before downstream usage. Samples with an average PFF size (Z-Ave) (Top Panel) of 50 nm +/-10 nm were employed in this study. Representative size distribution of PFFs in solution post-sonication can be seen in bottom Intensity (%) vs size (nm) graph. The target 50 nm peak is depicted by the red line.

**Supplemental Figure 2: Secondary Antibody Only Microscopy Controls for Pathological α-Synuclein**. A) Murine PFF injected cortical tissue no primary antibody staining in red channel at 60x magnification (diffraction limited) to show extent of off target staining (left). Human Synucleinopathy no primary control staining for red channel. Substantial autofluorescence and cellular debris staining can be observed on SORA super-resolution field of view.

**Supplemental Figure 3: Fluorescent Signal is converted to 3D Surface Rendering for Analysis. A)** Please see Supplemental Video 5 for animation depicting murine a-syn pathology from 3D fluorescence view to surface creation. Surfaces were created off of fluorescent signal to aid in analysis. **B)** Please see Supplemental Video 6 for human a-syn pathology animation in 3D fluorescence view and surfaces. This is performed on representative Lewy Body from Fig. 2D/F. Human sphericity quantification based on Lamina morphology (Right). Variance in nuclear integrity due to PMI was high within and between tissue (Right). Tissue 3 had more degradation, picked up more objects in same field of view, all of which prevented meaningful Lamina analysis with current workflow.

**Supplemental Figure 4**: **Lateral and Axial Resolution of Micrographs. A)** Axial Resolution of images was dependent on 300 nm step size for z-stack acquisition. 600 nm depth of p-α-syn penetration within the nucleus was used as a cutoff value for nuclear localization in all analysis to cautiously account for 300 nm resolution error in overlapping surfaces. Example measurement (human tissue) of two objects (left) shows clear distinction of objects at this resolution. The SoRa microscope achieves lateral resolution of 120 nm through optical reassignment. Therefore, a cutoff of 240 nm was used for lateral nuclear localization within the nuclear structure. Example measurement (Right) of two distinct objects in lateral view at roughly 120 nm resolution. **B)** Example axial measurement of mouse p-α-syn penetration within nuclear surfaces. Nuclear surfaces were made transparent to visualize p-α-syn within.

**Supplemental Figure 5**: **Human α-Syn Pathology Varies in Size and Morphologies. A)** Small p-α-syn aggregates assume various morphologies similar in structure to murine p-α-syn aggregates. In the merged image (Right) the yellow arrow points to a p-α-syn structure resembling murine Type II basket-like pathology. The green arrows point to a p-α-syn structure resembling Type I murine pathology classification.

**Supplemental Figure 6: Neuro 2a cells accumulate pathological a-Syn in nuclear compartments**. Addition of PFF’s to Neuro 2a cell cultures creates robust nuclear α-synuclein pathology. **A)** Immunofluorescence micrograph α-syn staining in Neuro 2a cells. Corrupted p-α-syn (Middle Image) is largely nuclear localized, while total α-syn immunofluorescence labeling, seen in green channel as small puncta (2^nd^ Image), has cell-filling distribution.

