## Supplemental Figures for "Pathological α-Synuclein Perturbs Nuclear Integrity"

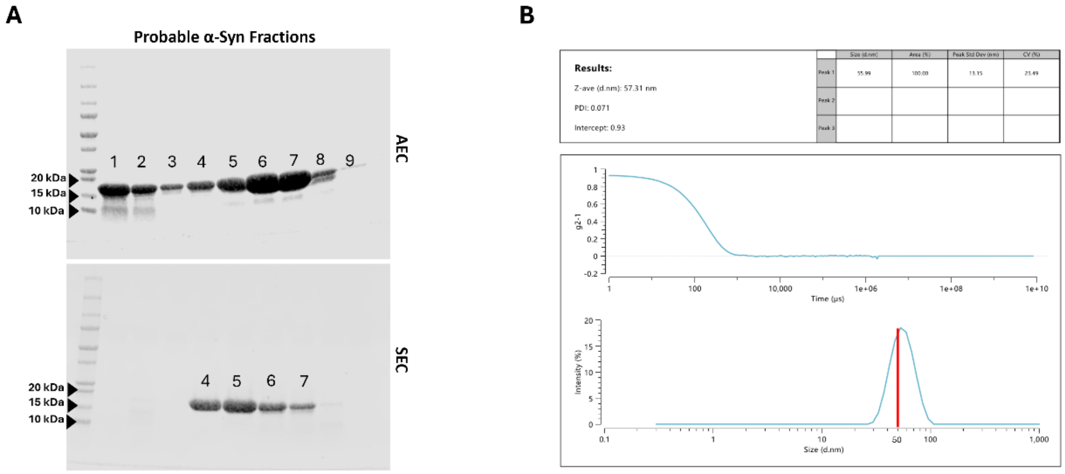


**Supplemental Figure 1: Quality Assurance Metrics for α-Synuclein Purification from BLD21 cells expressing PRK172 plasmid. A)** Gelcode Blue stained SDS-PAGE gels loaded with probable fractions generated from liquid chromatography. Top image depicts fractions after anion exchange (AEC), with nine total positive for α-syn. Bottom image depicts probable fractions after size exclusion chromatography (SEC), with four fractions positive for α-syn (fractions 4-7). **B)** Representative dynamic light scattering (DLS) sizing quality check of PFF samples before downstream usage. Samples with an average PFF size (Z-Ave) (Top Panel) of 50 nm +/- 10 nm were employed in this study. Representative size distribution of PFFs in solution post-sonication can be seen in bottom Intensity (%) vs size (nm) graph. The target 50 nm peak is depicted by the red line.

A

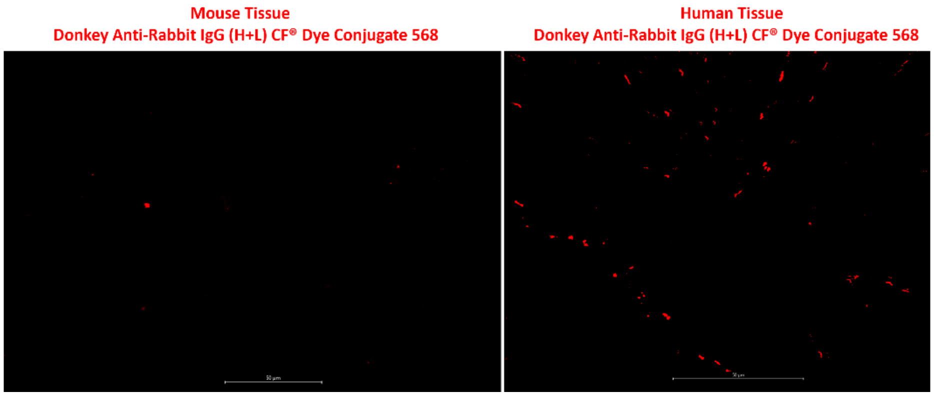


**Supplemental Figure 2: Secondary Antibody Only Microscopy Controls for Pathological α-Synuclein. A)** Murine PFF injected cortical tissue no primary antibody staining in red channel at 60x magnification (diffraction limited) to show extent of off target staining (left). Human Synucleinopathy no primary control staining for red channel. Substantial autofluorescence and cellular debris staining can be observed on SORA super-resolution field of view.

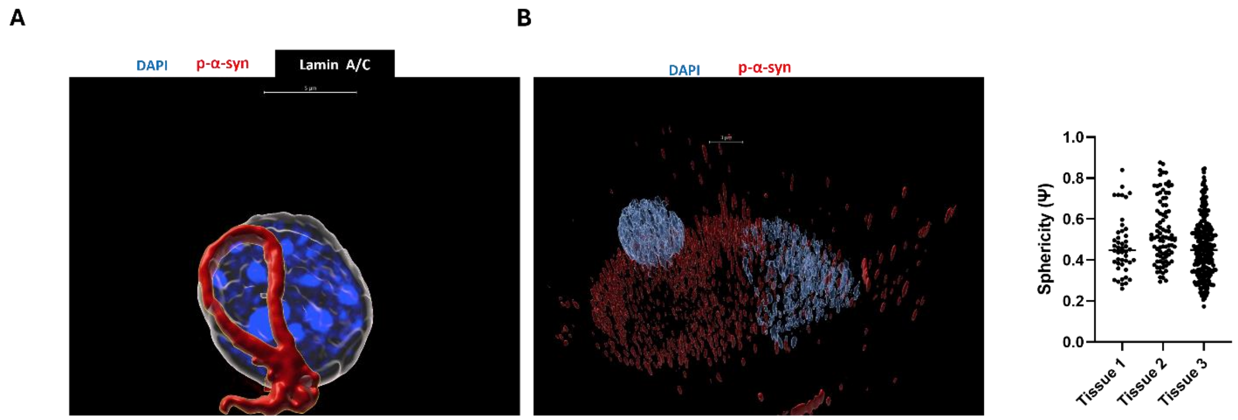


**Supplemental Figure 3: Fluorescent Signal is converted to 3D Surface Rendering for Analysis. A**) Please see Supplemental Video 5 for animation depicting murine a-syn pathology from 3D fluorescence view to surface creation. Surfaces were created off of fluorescent signal to aid in analysis. **B)** Please see Supplemental Video 6 for human a-syn pathology animation in 3D fluorescence view and surfaces. This is performed on representative Lewy Body from Fig. 2D/F. Human sphericity quantification based on Lamina morphology (Right). Variance in nuclear integrity due to PMI was high within and between tissue (Right). Tissue 3 had more degradation, picked up more objects in same field of view, all of which prevented meaningful Lamina analysis with current workflow.

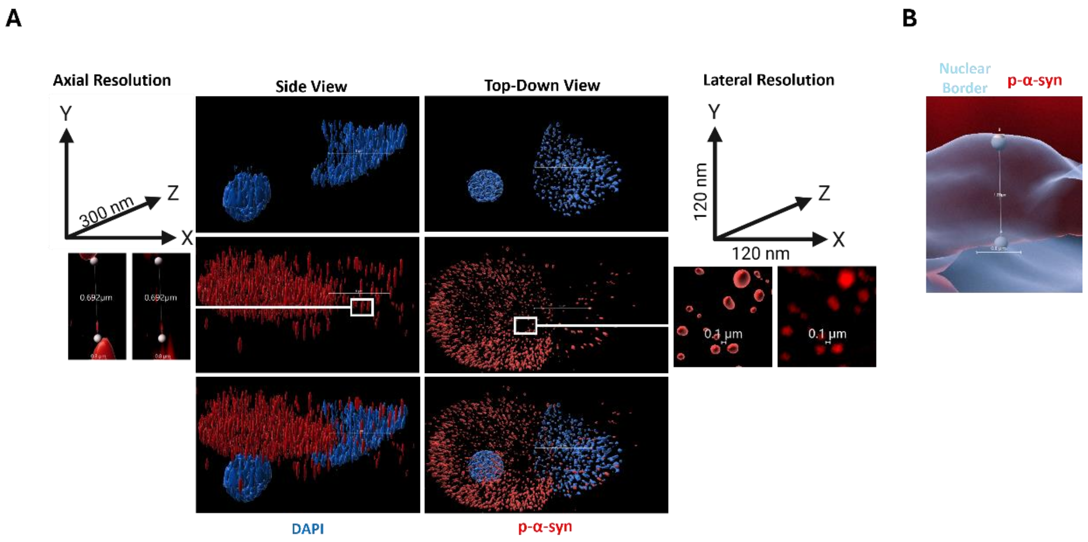


**Supplemental Figure 4: Lateral and Axial Resolution of Micrographs**. **A)** Axial Resolution of images was dependent on 300 nm step size for z-stack acquisition. 600 nm depth of p-α-syn penetration within the nucleus was used as a cutoff value for nuclear localization in all analysis to cautiously account for 300 nm resolution error in overlapping surfaces. Example measurement (human tissue) of two objects (left) shows clear distinction of objects at this resolution. The SoRa microscope achieves lateral resolution of 120 nm through optical reassignment. Therefore, a cutoff of 240 nm was used for lateral nuclear localization within the nuclear structure. Example measurement (Right) of two distinct objects in lateral view at roughly 120 nm resolution. **B)** Example axial measurement of mouse p-α-syn penetration within nuclear surfaces. Nuclear surfaces were made transparent to visualize p-α-syn within.

A

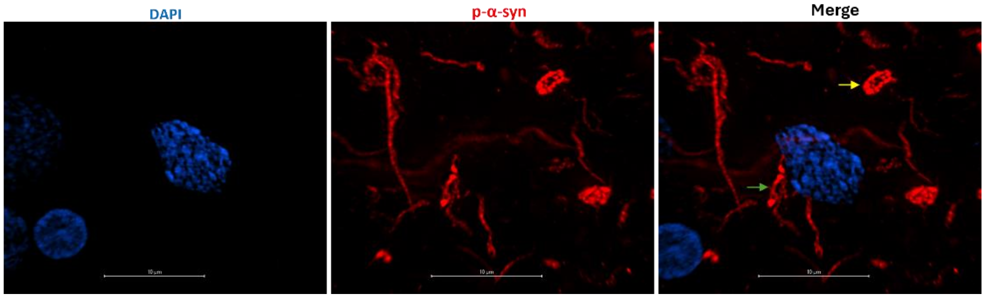


**Supplemental Figure 5: Human α-Syn Pathology Varies in Size and Morphologies. A)** Small p-α-syn aggregates assume various morphologies similar in structure to murine p-α-syn aggregates. In the merged image (Right) the yellow arrow points to a p-α-syn structure resembling murine Type II basket-like pathology. The green arrows point to a p-α-syn structure resembling Type I murine pathology classification.

A

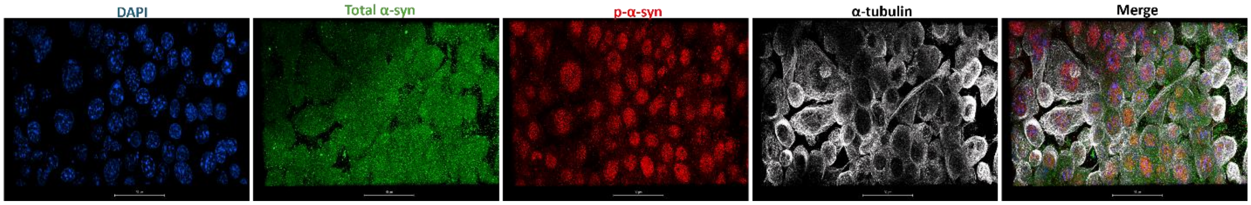


**Supplemental Figure 6**: **Neuro 2a cells accumulate pathological a-Syn in nuclear compartments.** Addition of PFF's to Neuro 2a cell cultures creates robust nuclear α-synuclein pathology. **A)** Immunofluorescence micrograph α-syn staining in Neuro 2a cells. Corrupted p-α-syn (Middle Image) is largely nuclear localized, while total α-syn immunofluorescence labeling, seen in green channel as small puncta (2nd Image), has cell-filling distribution.
